# A bimodular PKS platform that expands the biological design space

**DOI:** 10.1101/2020.05.14.096743

**Authors:** Amin Zargar, Luis Valencia, Jessica Wang, Ravi Lal, Samantha Chang, Miranda Werts, Andrew R. Wong, Veronica Benites, Edward Baidoo, Leonard Katz, Jay D. Keasling

## Abstract

Traditionally engineered to produce novel bioactive molecules, Type I modular polyketide synthases (PKSs) could be engineered as a new biosynthetic platform for the production of *de novo* fuels, commodity chemicals, and specialty chemicals. Previously, our investigations manipulated the first module of the lipomycin PKS to produce short chain ketones, 3-hydroxy acids, and saturated, branched carboxylic acids. Building upon this work, we have expanded to multi-modular systems by engineering the first two modules of lipomycin to generate unnatural polyketides as potential biofuels and specialty chemicals in *Streptomyces albus*. First, we produce 20.6 mg/L of the ethyl ketone, 4,6 dimethylheptanone through a reductive loop exchange in LipPKS1 and a ketoreductase knockouts in LipPKS2. We then show that an AT swap in LipPKS1 and a reductive loop exchange in LipPKS2 can produce the potential fragrance 3-isopropyl-6-methyltetrahydropyranone. Highlighting the challenge of maintaining product fidelity, in both bimodular systems we observed side products from premature hydrolysis in the engineered first module and stalled dehydration in reductive loop exchanges. Collectively, our work expands the biological design space and moves the field closer to the production of “designer” biomolecules.

**Highlights:**

- Engineered lipomycin module 1 and module 2 to produce unnatural polyketides as valuable bio-based chemicals
- A reductive loop swap and ketoreductase knockout used to produce 20 mg/mL of a novel ethyl ketone, a gasoline replacement
- An acyltransferase swap and reductive loop swap successfully produced δ-lactone, a potential fragrant compound
- Incomplete reduction and premature hydrolysis observed in engineered modules

**Graphical abstract:** 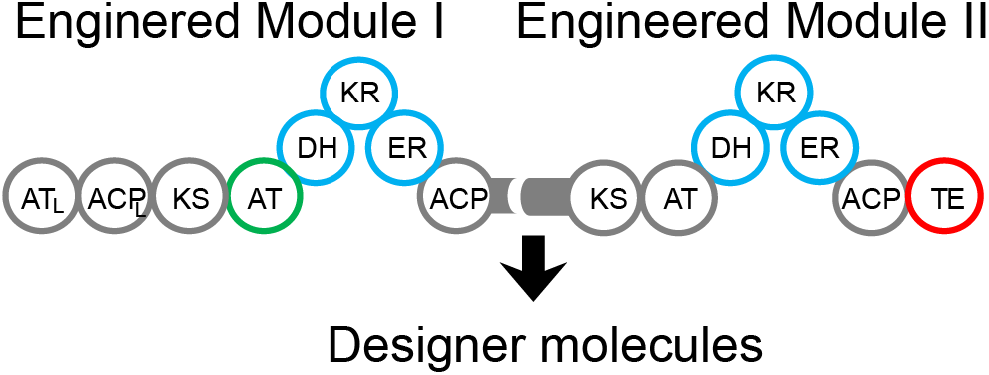

## 1. Introduction

Advances in biotechnology have only begun to capitalize on the biomolecular design space. Referred to as the parvome, ‘parv-’ meaning small and ‘-ome’ denoting group, the world of cell-based molecules is vastly larger than the known chemical design space (Davies, 2011). *De novo* biomolecular production efforts have sought to capitalize on this space to generate new biofuels, commodity chemicals, and specialty chemicals (King et al., 2016). Beyond developing molecules with superior properties, biosustainable production of these molecules could contribute to a substantial reduction in carbon emissions, which is needed to avoid potentially devastating climate change (Matthews et al., 2009). Generally, biosynthesis of unnatural molecules often relies on broad substrate ranges (Rodriguez et al., 2014) and promiscuous activity in enzymes (Khersonsky et al., 2006). While major advances have been made in protein engineering, redesigning proteins to generate novel bioactivity and achieve new products remains a major challenge (Kumar et al., 2018).

Polyketide synthases synthesize an astonishing diversity of natural products including, but not limited to, anticancer, antimicrobial, and immunomodulating compounds (reviewed by (Robbins et al., 2016)). Assembly-line, modular polyketide synthases (PKSs), a subset of Type I PKSs, are often linked in a collinear fashion, creating a design space that could be rationally reprogrammed to produce many valuable biomolecules (Cai and Zhang, 2018; Yuzawa et al., 2018b; Zargar et al., 2017, 2018). Each module’s cycle begins with a Claisen-like condensation reaction between the growing chain on the ketosynthase (KS) domain and a malonyl-CoA analog on the acyl carrier protein (ACP) that was loaded by the acyltransferase domain (AT) (**Figure 1A**). Unlike fatty acid synthases that exclusively incorporate malonyl-CoA, AT domains of Type-I PKSs select a wide variety of extender units, greatly expanding the biological design space. After chain extension, the molecule’s carbonyl reduction state is determined by the reductive domains present within a module, namely the ketoreductase (KR), dehydratase (DH), and enoylreductase (ER), which generate the β-hydroxyl, a-β alkene (typically *trans),* or saturated β-carbons respectively when progressively combined; PKSs can have variability in β-carbon reduction, which is a major source of polyketide diversity and another attractive feature for molecular design. Finally, a thioesterase (TE) domain typically releases the final product from the megasynthase via hydrolysis or cyclization. While these biomolecular pathways have most often been engineered to fine-tune potential drug candidates, combinatorial biosynthesis could be implemented to generate molecules with simple scaffolds, such as biofuels and industrial chemicals (Cai and Zhang, 2018). While combinatorial biosynthesis of PKSs through domain modification, module swaps, and other techniques have made major progress in drug development (Hertweck, 2015; Weissman, 2016; Wong and Khosla, 2012), *de novo* biomolecular production is still a nascent field, and there have been no examples of multi-modular PKS redesign to produce valuable biochemicals.

**Figure 1.**
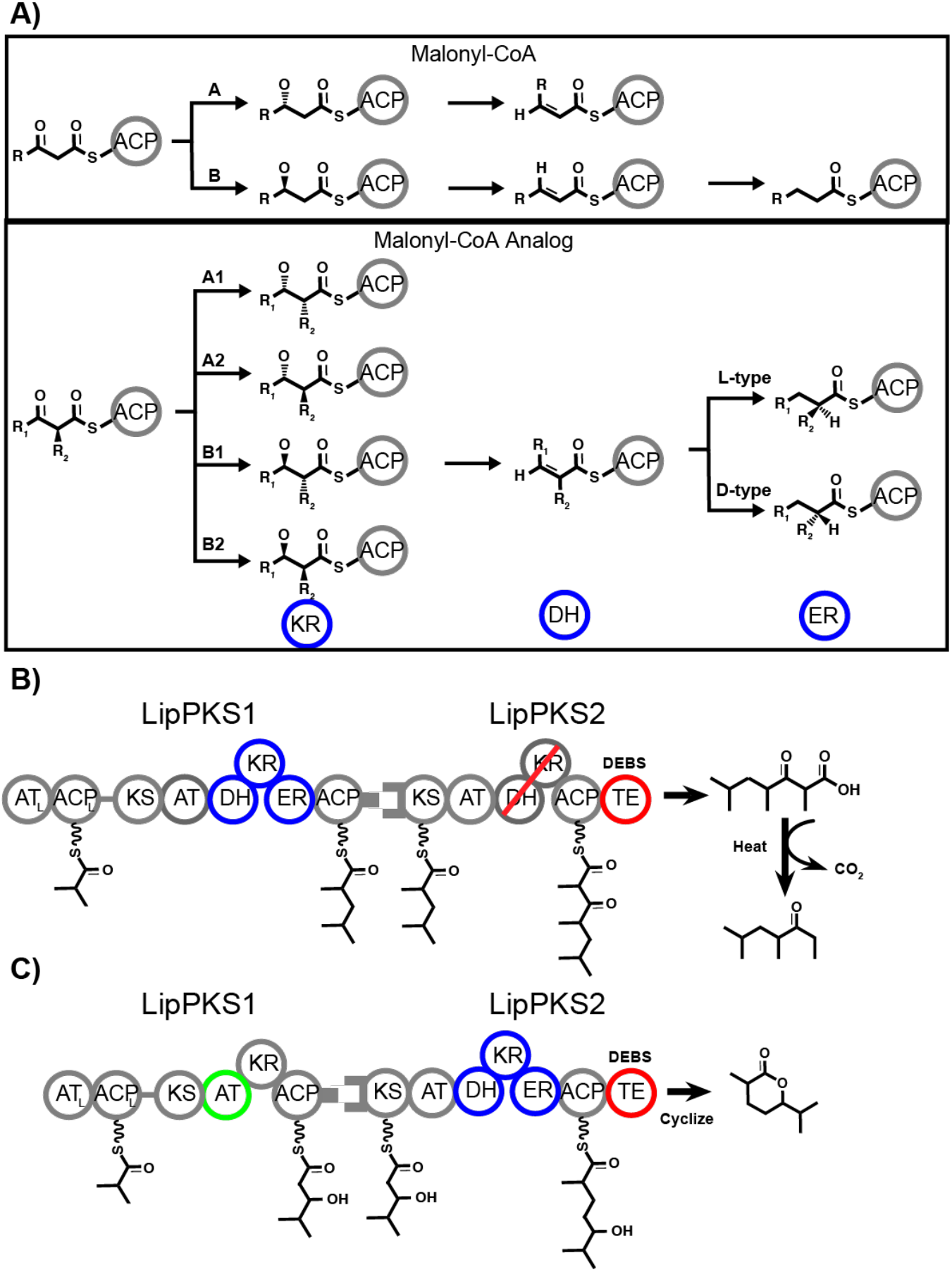
Schematic of PKS processing and engineering design in this study. **A)** PKS processing of each subtype of malonyl-CoA and malonyl-CoA analog extender units **B)** Lipomycin bimodular PKS design to produce ethyl ketones through a full reductive donor loop in LipPKS1 (blue circles), a KR mutant to abolish activity (red line), and a fused DEBS TE (red circle) **C)** Lipomycin bimodular PKS design to produce δ-lactone through an AT-swap in LipPKS1 (green circle), a full reductive donor loop in LipPKS2 (blue circles), and a fused DEBS TE (red circle).

Previously, our group has engineered three major PKS elements in the first module of lipomycin: 1) an inserted TE to produce 3-hydroxy acids (Yuzawa et al., 2017a), 2) a KR knockout and AT domain swap to produce short chain ketones (Yuzawa et al., 2018a, 2017b), and 3) reductive loop (RL) exchanges to produce saturated, short chain carboxylic acids (Zargar et al., 2019). The design space expands considerably with multiple module systems, and in this work, we build on our single module platform by combining multiple PKS manipulations (KR knockouts, reductive loop (RL) swaps, AT swaps, fused TE) in a biomodular system to produce novel biomolecules, namely biofuels and specialty chemicals.

## 2. Materials and Methods

### 2.1 Cloning

#### Cloning of all constructs

All clusters were expressed from the *Streptomyces albus* genome under control of the GapDH(El) promoter from *Eggerthella lenta.* Junction sites for reductive loop exchanges were determined by those reported by Hagen *et al.* through multiple sequence alignment with MUSCLE (Hagen et al. 2016; Edgar 2004). The plasmids along with their associated information have been deposited in the public version of JBEI registry (http://public-registry.jbei.org) and are physically available from the authors upon request https://public-registry.jbei.org/folders/557.

#### Cloning of LipPKS1 with full reductive loop modules and native docking domain

The φC31-based *Streptomyces* integrase vectors were used as described by Phelan *et al* to integrate the LipPKS1 reductive loop swap modules (Phelan et al., 2017). The native docking domain sequences of LipPKS1 were codon optimized for *E. coli* and synthesized by Gen9 (since acquired by Ginkgo Bioworks). They were cloned through Golden Gate assembly into the LipPKS1 module with an inserted RL from NanA2 from Zargar *et al* (Zargar et al., 2019).

#### Cloning of LipPKS1 with AT-swap and native docking domain

The φC31-based integrase vectors (Phelan et al., 2017) were used to integrate the AT-swapped LipPKS1 module into the φC31 site in the *S. albus* chromosome. The Lip1 native C-terminal docking domain sequences of LipPKS1 were cloned into the AT-swapped LipPKS1 module to replace the DEBS TE from Yuzawa *et al.* (Yuzawa et al., 2017b) through Golden Gate assembly.

#### Cloning of LipPKS2 with KR knockout and fused DEBS thioesterase

The VWB-based *Streptomyces* integrase vectors were used to integrate the LipPKS2 KR-module (Phelan et al., 2017). The native LipPKS2 was codon optimized for *E. coli* with a single point mutation S1547A into the KR active site to mutate the catalytic serine to alanine, thereby abolishing KR activity, and synthesized by Gen9 (since acquired by Ginkgo Bioworks). The fused DEBS thioesterase domain was placed at the C-terminus of the ACP domain through Golden Gate assembly.

#### Cloning of LipPKS2 with full reductive loop modules and fused DEBS thioesterase

The VWB *Streptomyces* integrase vectors were used to integrate the LipPKS2 reductive loop modules cloned previously (Zargar et al., 2019).

### 2.2 Genetic integration

#### Conjugation of LipPKS1 constructs into *S. albus* J1074

*Escherichia coli* ET12567/pUZ8002 was transformed with LipPKS1 plasmids and selected for on LB agar containing kanamycin (25 μg/mL), chloramphenicol (15 μg/mL), and apramycin (50 μg/mL). A single colony was inoculated into 5 mL of LB containing kanamycin (25 μg/mL), chloramphenicol (15 μg/mL), and apramycin (50 μg/mL) at 37°C. The overnight culture was used to seed 10 mL of LB containing the same antibiotics, and the new culture was grown at 37°C to an OD600 of 0.4-0.6. The *E. coli* cells were pelleted by centrifugation, washed twice with LB, and resuspended in 500 μL of LB. Fresh *S. albus* J1074 spores were collected from a mannitol soy agar plate with 5 mL of 2xYT and incubated at 50°C for 10 min. The spores (500 μL) and the *E. coli* cells (500 μL) were mixed, spread onto mannitol soy agar, and incubated at 30°C for 16 hours. 1 mL of both nalidixic acid (20 μg/mL) and apramycin (40 μg/mL) were added to the plate and allowed to dry.The plate was then incubated for 3-4 days at 30°C.

*A* single colony was inoculated into TSB containing nalidixic acid (25 μg/mL) and apramycin (25 μg/mL). After 3-4 days, a 1 mL aliquot was taken for genomic isolation using the Maxwell kit (Promega, Cat# AS1490, Madison WI). Successful integration was verified using qPCR. The remainder of the culture was spread onto a MS plate and incubated at 30oC for 2-3 days. The spores were collected from the plate with 3-4 mL of water and mixed with glycerol to prepare a 25% glycerol stock, which was stored at −80°C for long-term storage.

#### Conjugation of VWB integrase LipPKS2 constructs into LipPKS1-conjugated *S. albus*

*E. coli* ET12567/pUZ8002 was transformed with LipPKS2 plasmids and selected for on LB agar containing kanamycin (25 μg/mL), chloramphenicol (15 μg/mL), and spectinomycin (200 μg/mL). A single colony was inoculated into 5 mL of LB containing kanamycin (25 μg/mL), chloramphenicol (15 μg/mL), and spectinomycin (200 μg/mL) at 37°C. The overnight culture was used to seed 10 mL of LB containing the same antibiotics, and the new culture was grown at 37°C to an OD600 of 0.4-0.6. The *E. coli* cells were pelleted by centrifugation, washed twice with LB, and resuspended in 500 μL of LB. *S. albus* spores with an integrated LipPKS1, which were collected from a mannitol soy agar plate with 5 mL of 2xYT and incubated at 50°C for 10 min. The spores (500 μL) and the *E. coli* cells (500 μL) were mixed, spread onto mannitol soy agar, and incubated at 30°C for 16 hours. 1 mL of each nalidixic acid (20 μg/mL), apramycin (40 μg/mL), and spectinomycin (400 μg/mL) was added to the plate and allowed to dry. The plate was then incubated for 3-4 days at 30°C. A single colony was inoculated into TSB containing nalidixic acid (25 μg/mL), apramycin (25 μg/mL) and spectinomycin (200 μg/mL). After 3-4 days, a 1 mL aliquot was taken for genomic isolation through the Maxwell kit (Promega, Cat# AS1490, Madison WI). Successful integration was verified through qPCR. The remainder of the culture was spread onto a MS plate and incubated at 30oC for 2-3 days. The spores were collected from the plate with 3-4 mL of water and mixed with glycerol to prepare a 25% glycerol stock, which was stored at −80°C for long-term storage.

### 2.3 Production runs

#### *S. albus* production runs

Engineered *S. albus* spores were grown in 12 mL of TSB medium containing nalidixic acid (50 μg/mL), apramycin (50 μg/mL) and spectinomycin (200 μg/mL) for 4-5 days at 30°C. Three mL of the overnight culture was used to seed 30 mL of 10% media 042 and 90% plant hydrolysate (Yuzawa et al., 2018a), supplemented with 2.4 grams/liter of valine and nalidixic acid (50 μg/mL), which was grown for 10 days at 30°C. For production runs of lactones, an overlay of 4 mL of dodecane was added to retain the product.

### 2.4 Sample preparation

#### Sample preparation for detection of acids

To detect acid side products, 1 mL of each sample was centrifuged at 5000 g for 10 minutes and 200 μL of the supernatant was removed. The supernatant was mixed with 200 μl of 100 μM hexanoic acid dissolved in methanol and filtered using Amicon Ultra Centrifugal filters, 3 KDa Ultracel, 0.5 mL device (Millipore). β-hydroxy (3-hydroxy-2,4-dimethylpentanoic acid) and saturated acids (2,4-dimethylpentanoic acid) were synthesized by Enamine (Cincinnati, USA) to greater than 95% purity.

#### Sample preparation for bimodular production of triketide lactones

10 mL of each sample was mixed with 2 mL of diethyl ether in a 15-mL conical tube and vortexed for 5 minutes. Each conical tube was centrifuged at 5000g for 10 minutes and 1 mL of ether was removed and placed into a 2-mL flat bottom microcentrifuge tube. Air was gently blown over each sample in a chemical fume hood until dry. The extract was resuspended in 200 μL of methanol. 5-methyl-6-(propan-2-yl)oxan-2-one was synthesized by Enamine (Cincinnati, USA) to greater than 95% purity.

#### Sample preparation for detection of 4,6-dimethyl heptanone

One mL of each sample was harvested in a 1.7-mL microcentrifuge tube. To each tube, 300 μL of ethyl acetate and 50 μL of formic acid were added. All tubes were wrapped in paraffin and heated for 60 minutes at 80C. Samples were then placed on ice for 5 minutes and vortexed for 5 minutes. Each sample was centrifuged for 2 minutes at 10,000g. One hundred microliters of ethyl acetate was removed from each sample and placed in a GC MS vial.

### 2.5 Analytical chemistry

#### GC-MS detection of 4,6-dimethyl heptanone

Electron ionization GC/MS analysis was performed on a G3950A-9000 GC (Agilent) using a J&W HP-5ms Ultra Inert Intuvo GC column module (15 m length, 0.25 mm inner diameter, 0.25 μm film thickness). The GC was coupled to a mass selective detector (Agilent 5977B MSD) and an autosampler (Model 7693 Agilent). The GC oven was programmed at 60°C for 3 minutes, ramping at 10°C/ min until 120°C, and then ramping at 200°C/min to 300°C; the injection port temperature was 250°C. Using an authentic standard, we determined a single-ion method of detection collecting data at m/z = 57.00, m/z=85.00, m/z=142.00.

#### LC-MS detection of short chain acids

The LC-MS analysis was conducted on a Kinetex XB-C18 column (100-mm length, 3.0-mm internal diameter, and 2.6-μm particle size; Phenomenex, Torrance, CA USA) using an Agilent Technologies 1200 Series HPLC system. The mobile phase for separating 2,4-dimethylpentanoic acid and 2,4-dimethylpent-2-enoic acid was composed of 10 mM ammonium acetate and 0.05% ammonium hydroxide in water (solvent A) and 10 mM ammonium acetate and 0.05% ammonium hydroxide in methanol (solvent B). The mobile phase for separating 3-hydroxy-2,4-dimethylpentanoic acid and 2,3-dimethyl-3-oxopentanoic acid was composed of 0.1% formic acid in water (solvent A) and 0.1% formic acid in methanol (solvent B). All acids were each separated via the following gradient: increased from 5 to 97.1% B in 6.5 min, held at 97.1% B for 1.3 min, decreased from 97.1 to 5% B in 0.4 min, and held at 5% B for an additional 2 min. The flow rate was held at 0.42 ml · min-1 for 8.2 min, and then increased from 0.42 to 0.65 ml · min-1 for an additional 2 min. The total LC run time was 10.8 min. Samples of 3 μl were injected into the LC column. Acids were detected via [M-H]-ions. Nitrogen gas was used as both the nebulizing and drying gas to facilitate the production of gas-phase ions. The drying and nebulizing gases were set to 11 L · min^-1^ and 30 L · bin^-2^, respectively, and a drying gas temperature of 330°C was used throughout. Atmospheric pressure chemical ionization was conducted in the positive-ion mode with capillary and fragmentor voltages of 3.5 kV and 100 V, respectively. The skimmer, OCT 1 RF, and corona needle were set to 50 V, 170 V, and 4 μA, respectively. The vaporizer was set to 350°C. The analysis was performed using an *m/z* range of 70 to 1100. Data acquisition and processing were performed using MassHunter software (Agilent Technologies, United States).

#### LC-MS detection of triketide lactones

LC separation of triketide lactones was conducted on a Kinetex XB-C18 reversed phase column (100 mm length, 3 mm internal diameter, 2.6 μm particle size; Phenomenex, United States) using an Agilent 1200 Rapid Resolution LC system (Agilent Technologies, United States). The mobile phase was composed of water (solvent A) and methanol (solvent B). Lactones were each separated via the following gradient: increased from 30 to 90% B in 3.7 min, held at 94% B for 5.2 min, decreased from 90 to 30% B in 0.33 min, and held at 30% B for an additional 2.0 min. The flow rate was held at 0.42 ml · min^-1^ for 8.67 min, increased from 0.42 to 0.60 ml · min^-1^ in 0.33 min, and held at 0.60 ml · min^-1^ for an additional 2.0 min. The total LC run time was 11.0 min. The column compartment and autosampler temperatures were set to 50°C and 6°C, respectively. Samples of 3 μL were injected into the LC column. The Agilent 1200 Rapid Resolution LC system was coupled to an Agilent 6210 TOF (Agilent Technologies, United States). Nitrogen gas was used as both the nebulizing and drying gas to facilitate the production of gas-phase ions. The drying and nebulizing gases were set to 10 L · min^-1^ and 25 L · bin, respectively, and a drying gas temperature of 325°C was used throughout. Atmospheric pressure chemical ionization was conducted in the positive-ion mode with capillary and fragmentor voltages of 3.5 kV and 100 V, respectively. The skimmer, OCT1 RF, and corona needle were set to 50 V, 170 V, and 4 μA, respectively. The vaporizer was set to 350°C. The analysis was performed using an *m/z* range of 70 to 1100. Data acquisition and processing were performed using MassHunter software (Agilent Technologies, United States).

## 3.0 Results

### 3.1 Production of ethyl ketones

Short chain ketones (C3-C7) have been noted for their potential as gasoline additives because of their high octane numbers (McCormick et al., 2017). We recently tested their fuel combustion properties in a gasoline called CARBOB, a specially formulated Blendstock for Oxygenate Blending formula mandated by the state of California (Yuzawa et al., 2018a). While most ketones showed superior properties to the common biofuel butanol (octane numbers, energy density, boiling point, melting point, and flash point), methyl-branched C5 and C6 ketones had comparable fuel properties to isooctane. Longer chain ketones (above C7) were too expensive to synthesize for testing, but it is likely they are candidates as gasoline blending agents as well, and possibly possess combustion properties comparable or superior to traditional gasoline molecules. We therefore sought to produce longer chain ketones as an example of *de novo* production of biomolecules in a bimodular PKS system.

As illustrated in **Figure 2A**, we aimed to produce ethyl ketones through an RL exchange in LipPKS1 and an AT swap in LipPKS2. Previously, we had performed an RL exchange in LipPKS1 with a DEBS1-TE, and found a correlation between successful production of the desired product and the chemical similarity of the donor and recipient reductive loops with the most production through the chimera of LipPKS1 with an inserted donor loop form NanA2 (nanchangamycin, module 2) (Zargar et al., 2019). In the work presented here, we replaced the fused DEBS (6-deoxyerythronolide) thioesterase with the native docking domain of LipPKS1. For LipPKS2, we synthesized the codon-optimized gene with the native docking domain and a single point mutation at S1547A to mutate the catalytic serine to alanine in the KR domain. With the thioesterase of DEBS inserted following the ACP domain of LipPKS2, the programmed product of engineered Lip1 - Lip2 is a β-keto carboxylic acid, which, upon acidification and heat, is an ethyl ketone, 4,6-dimethyl heptanone.

**Figure 2.**
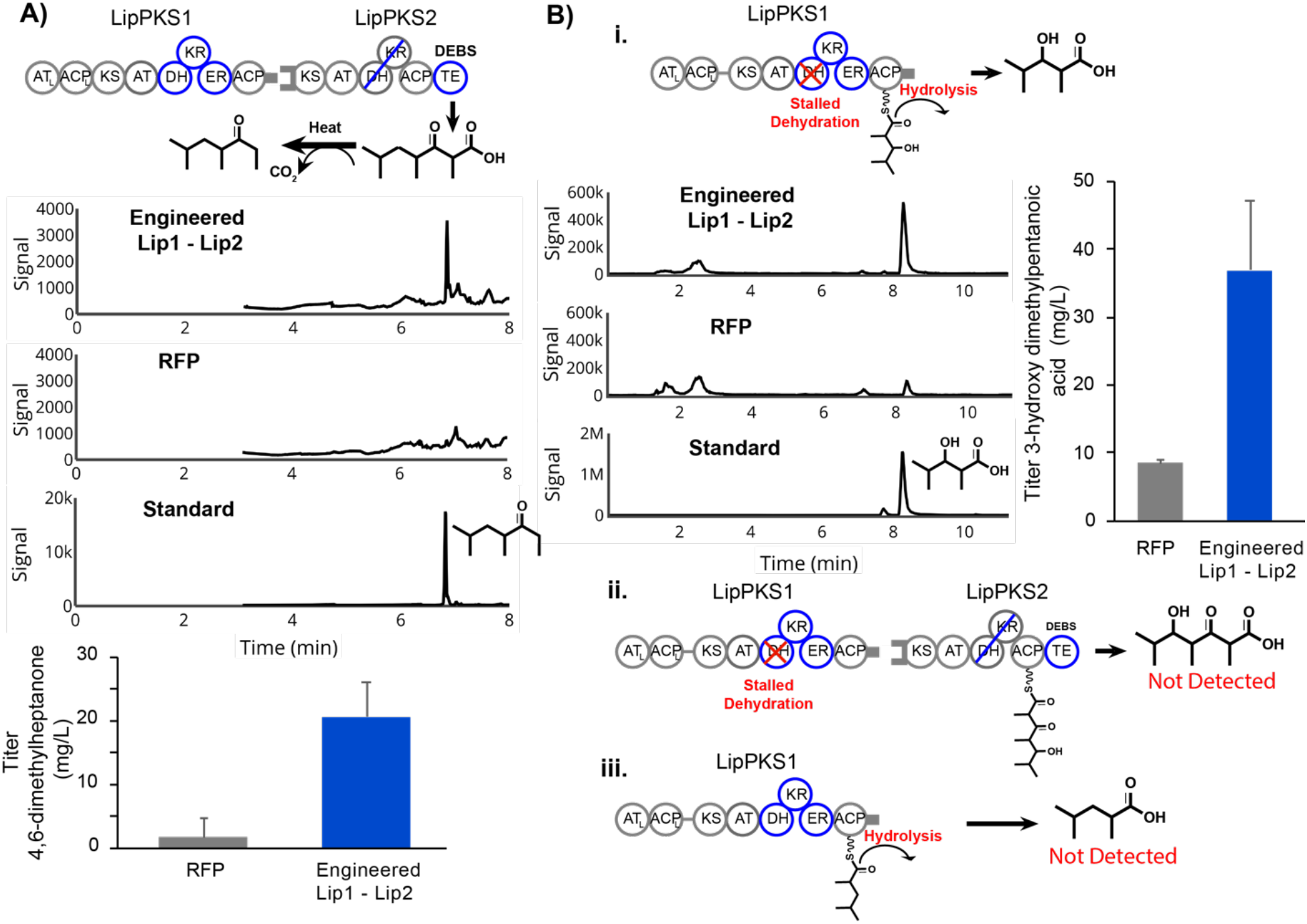
Production of ethyl ketones and side products in engineered Lip1 - Lip2 bimodular system. A) Schematic, MS chromatogram, and quantification of 4,6-dimethyl heptanone **B) i.** Schematic, MS chromatogram and quantification of the side product 3-hydroxy-2,4- dimethylpentanoic acid due to incomplete reduction by LipPKS1 **ii.** Schematic of the side product of incomplete reduction in LipPKS1 processed and elongated by LipPKS2 **iii.** Schematic of the side product of complete reduction in LipPKS1 with premature hydrolysis

To produce the ethyl ketones, we conjugated the engineered Lip1 and Lip2 with the φC31 or VWB integrases, respectively, into the genome of *S. albus* J1074. After 10-day production runs, we harvested the samples and measured titers of the final product and side products. We observe production of the desired product after heating and acidification with a titer of 20.6 mg/L (**Figure 2A**). While successful, this was a considerable drop in titer compared to the 165 mg/L of the saturated, carboxylic acid produced by the singular Lip1 extension module with NanA2 reductive loops and a fused TE. This loss of production is partially reflected in the amount of side products generated in the bimodular system. Previously, we found that the unimodular LipPKS with NanA2 reductive loops and a fused TE produced the DH-stalled product, 3-hydroxydimethylpentanoic acid (Zargar et al., 2019). We therefore suspected that incomplete β-carbon reduction by the engineered module 1 could cause premature hydrolysis of the product from the Lip1 ACP, resulting in production of the DH-stalled product, 3-hydroxy 2,4-dimethylpentanoic acid, which was produced at a titer of 36.9 mg/L, considerably higher than that produced by the negative control (**Figure 2Bi**). Previous studies on *cis*-AT PKS modules have shown that the elongating condensation reaction at the KS between the substrate and extender unit has higher selectivity than acylation of the KS by the substrate, a process generally known as gatekeeping (Watanabe et al., 2003; Wu et al., 2004). This may also explain the loss in titer compared to the engineered unimodular Lip1 (~165 mg/L), as the stalled KS could reduce turnover. However, we did not observe this stalled β-hydroxy compound passed onto LipPKS2 and processed by the second module (**Figure 2Bii**). Importantly, the native Lip1 KR and donor NanA2 RL loop produce β-hydroxy compounds with different stereochemistries (A2 type compared to B1, **Figure 1A**). As it has been shown that KR domain exchanges generally retain native stereospecificity (Kao et al. 1998), we hypothesize that this difference in stereochemistry likely causes the downstream Lip2 KS to fail to elongate the stalled B1 type β-hydroxy substrate of the engineered LipPKS1. On the other hand, we did not detect any of the fully reduced product, 2,4-dimethyl pentanoic acid, prematurely hydrolyzing as a saturated acid (**Figure 2Biii)**. This is in keeping with other studies that show KS domains have less promiscuity with bulkier substrates (Jenner, 2016).

### 3.2 Production of δ-lactones

Over the past ten years, the market for genetically modified microbial production of fragrance aroma chemicals has grown. An important fragrant compound class is δ-lactones (Gupta, 2015), and while not commercially used, 3-isopropyl-6-methyltetrahydropyranone has been synthesized previously as a potentially fragrant δ-lactone (Plessis and Derrer, 2001). In a bimodular system with native AT domains, we previously engineered LipPKS1 and LipPKS2 to produce 3-isopropyl-4,6-dimethyltetrahydropyranone (Zargar et al., 2019), and in this work, we sought to incorporate AT-engineering to produce 3-isopropyl-6-methyltetrahydropyranone as an initial proof-of-concept production into this broad category of fragrant molecules.

The engineered production of this compound requires a combination of an AT swap in LipPKS1 and a RL swap in LipPKS2. For the AT swap in LipPKS1, we had previously analyzed AT domains and associated linkers to identify the boundaries for AT swaps while maintaining enzyme activity in LipPKS1 (Yuzawa et al., 2017b). We found that swapping the native methylmalonyl-CoA-selecting AT of LipPKS1 for the malonyl-CoA-selecting AT of the first module of the borreledin PKS (BorAT1), along with a fused DEBS TE, would result in the engineered unimodular system to produce 3-hydroxy carboxylic acids, which could be used as organic building blocks (Yuzawa et al., 2017b). Here, we replaced the fused TE with the native Lip1 docking domains in the BorAT1-swapped LipPKS1. For LipPKS2, we had previously introduced the NanA2 reducing loop, and when combined with the native LipPKS1, we had engineered production of 3-isopropyl-4,6-dimethyltetrahydropyranone (Zargar et al., 2019). This combination of LipPKS1 with the AT swap and LipPKS2 with an RL exchange should produce 3-isopropyl-6-methyltetrahydropyranone (Fig. 2A).

As before, we integrated the genes encoding the engineered LipPKS1 and LipPKS2 into the genome S. albus J1074 at the iφC31 and VWB phage attachment sites with φC31 and VWB integrases, respectively. After 10 day production runs, we detected the programmed product with a titer of 40 μg/L (**Figure 3A**). We did not observe the product in the acid form, as 5-dihydroxy 2,6-dimethyl heptanoic acid. This titer is considerably lower than the ethyl ketone titers from a similar bimodular PKS (20 mg/L). This could be caused by several factors including the non-native substrate presented to LipPKS2 by LipPKS1 with the malonyl-AT swap, disruption of the docking domain by the RL swap in LipPKS2, and lower turnover through the RL swap in LipPKS2. Once again, manipulation of the first module resulted in side products generated by premature hydrolysis at the LipPKS1. Premature hydrolysis of the AT swapped LipPKS1 would produce 3-hydroxy 4-methyl pentanoic acid, and we determined peaks of that mass in the culture (**Figure 3Bi**). As expected, we also observed production of 3,5-dihydroxy 2,6-dimethyl heptanoic acid, as the KR domain of the NanA2 reductive loop within LipPKS2 did not fully dehydrate the intermediate (**Figure 3Bii**). The stalled DH product in the cyclized form, 5-hydroxy-3-isopropyl-6-methyltetrahydropyranone was not detected.

**Figure 3.**
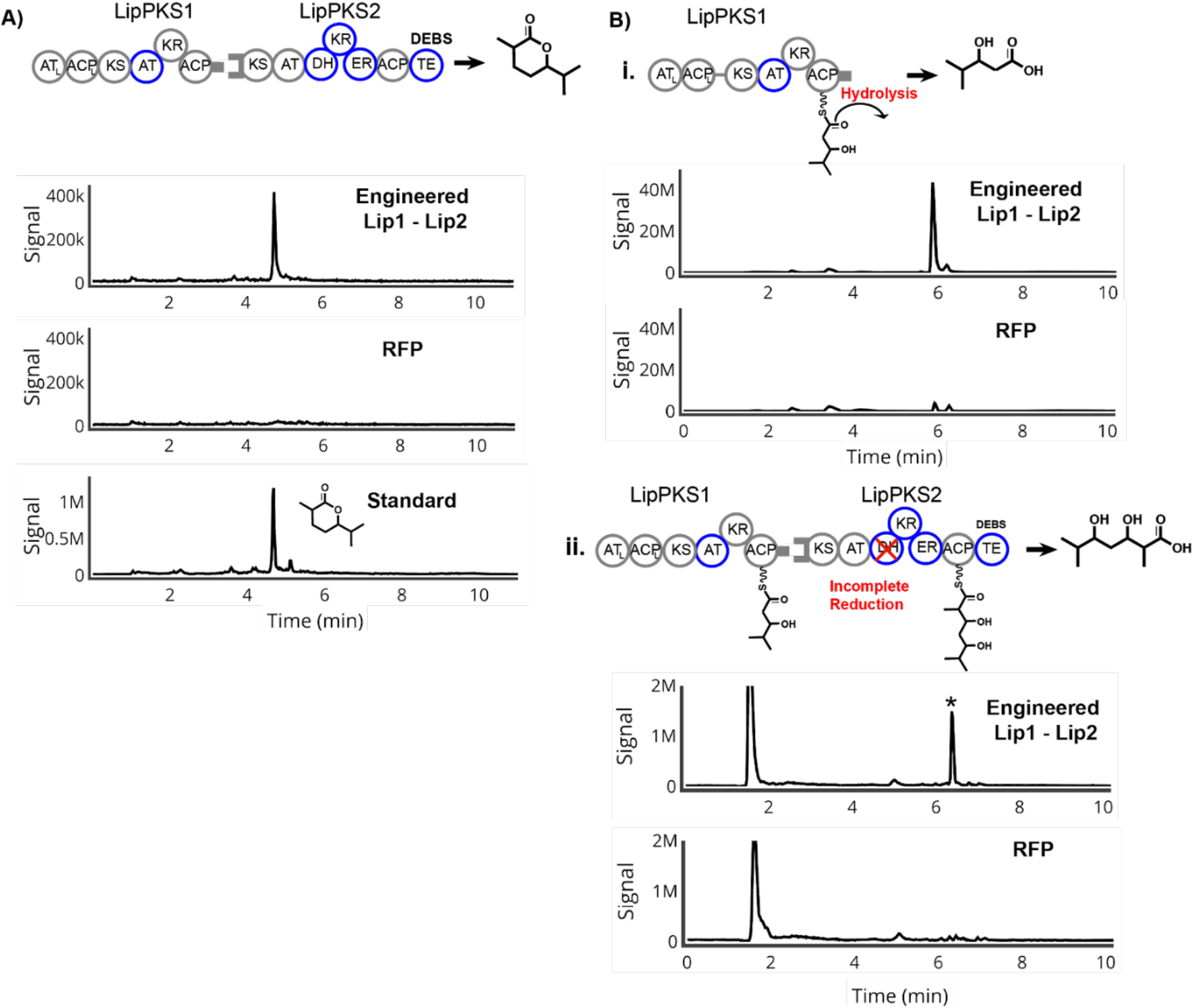
Production of δ-lactone and side products in engineered Lip1 - Lip2 bimodular system. **A)** Schematic, MS chromatogram, and quantification of 3-isopropyl-6-methyltetrahydropyranone **B) i.** Schematic and MS chromatogram of the side product 3-hydroxy-4-methylpentanoic acid due to premature hydrolysis of LipPKS1 **ii.** Schematic and MS chromatogram of the side product of incomplete reduction in LipPKS2

## Discussion

While unaltered natural products and synthetic chemistry have been the basis of industrial molecules, *de novo* biomolecular designs allow an astonishing diversity of molecules that could have a transformational impact in many fields (Smanski et al., 2016). A biosynthetic platform based on PKSs represents a vast design space with an attractive programming basis. With the diversity of starter substrates (~10^2^), malonyl-CoA analogs (~10^1^), and stereochemistry arrangements (~10^1^), a unimodular system alone can feasibly produce over 10,000 molecules and each subsequent module increases the number by two orders of magnitude. In the future, as our skills develop, we may be able to integrate our knowledge with other fields such as ‘click’ chemistry to obtain new capabilities (Le Feuvre and Scrutton 2018; Kalkreuter et al. 2019; Zhuet al. 2015). This would widely broaden both the scope of possible novel chemicals and our capacity to meet demands currently unable to be fulfilled.

In this study we’ve used a repertoire of AT domain swaps, mutagenesis, and reductive loop swaps with the goal of engineering a bimodular PKS while minimizing the disruption of protein-protein docking interactions. AT domain swaps enable the incorporation of rare extender units found in nature or even unnatural orthogonal extender units with click handles (Zhu et al. 2015) to expand the accessible chemical space. In our work, we performed an AT-swap to change the α-substituent of our growing polyketide chain with the principles described before to successfully change the extender unit of LipPKS1 from methylmalonyl-CoA to malonyl-CoA (Yuzawa et al. 2017). The other major factor in polyketide diversity is the degree of β-carbon reduction, particularly for biofuels where oxygenation can directly affect physical attributes such as melting temperature, hygroscopicity, H:C ratio, and vulnerability to oxidation (Wadumesthrige et al. 2009). We successfully employed a KR active site mutation in LipPKS2 to completely bypass reductive loop processing and ultimately yield ethyl ketones through the most conservative approach of active site mutagenesis. In contrast to the minimal disruption of a KR knockout, using reductive loop engineering principles recently described (Hagen et al. 2016) (Zargar et al. 2019), we were able to convert both LipPKS1 and LipPKS2 into fully reducing modules. This versatile bimodular platform paves the road for translating this strategy to a variety of PKSs with different loading modules and extension substrates to yield a diverse set of molecules. Additionally, a natural next step would involve extrapolating these engineering principles to PKS systems with more than 2 modules.

Failed production, reduced yield, and loss of product fidelity remain major challenges in PKS engineering. Online database tools such as ClusterCad (Eng et al., 2018) and SBSPKS (Khater et al. 2017) can facilitate PKS engineering, which can minimize the risk of failed production. A larger challenge is to increase fidelity and improve yield in multimodular systems,particularly as these systems are more likely to lose product fidelity as there is an added variable of intermodular interactions. Loss of product fidelity is not limited to domain and loop swaps as in this work; a recently engineered bimodular system successfully incorporated two non-native extender units through AT mutagenesis, but also produced unexpected side products through gatekeeping (Kalkreuter et al., 2019). Targeted mutagenesis of the ketosynthase can relieve the gatekeeping mechanisms, but the resulting activity is promiscuous, leading to loss of product fidelity (Jenner et al., 2015). Increased knowledge of PKS structure and function as well as machine learning algorithms on large datasets may inform future PKS designs to maximize product fidelity. Lastly, while improvements have been made in fermentation engineering to increase yield in hosts such as *S. albus,* strain engineering is still required to translate laboratory scale fermentation to industrial levels.

Previously, we have engineered the first module of lipomycin to make an array of molecules through AT swaps, RL swaps, KR knockouts, and a fused TE. Here, we have shown that we can leverage the knowledge we gained in unimodular systems to make multiple engineering manipulations to bimodular systems to produce novel biomolecules. While this is a step towards realizing the goals of capturing the biochemical design space, the unexpected metabolites produced through loss product fidelity highlight the need for a multi-factorial approach towards engineering these systems. Nonetheless, we successfully generated novel biofuels and specialty chemicals in the host *Streptomyces albus.*

## Acknowledgements

This work was funded by the DOE Joint BioEnergy Institute (http://www.jbei.org) supported by the U.S. Department of Energy, Office of Science, Office of Biological and Environmental Research, through contract DE-AC02-05CH11231 between Lawrence Berkeley National Laboratory and the U.S. Department of Energy, and the National Institute of Health Awards F32GM125179.

